# Dopamine controls the Sensitivity to Manganese induced Dopaminergic neurotoxicity in *Caenorhabditis elegans*

**DOI:** 10.1101/2024.10.23.619952

**Authors:** Vishnu Raj, Anoopkumar Thekkuveettil

## Abstract

Manganism, a disease with characteristic degeneration of dopamine neurons, has distinct aetiology and clinical manifestations, striking similarities with Parkinson’s Disease (PD). Environmental exposure to manganese (Mn) is one of the risk factors for the occurrence of PD. The definitive role of dopamine (DA) in Mn mediated neurodegeneration, and its developmental impact has not been well studied. To understand the pathways involved in Mn-induced neurotoxicity, we used *C. elegans* as the model system. Our results showed that adult worms treated with 50 and 100 mM MnCl_2_ significantly increased DA neurodegeneration. L1 larvae spared without neurodegeneration when treated with MnCl_2_ alone showed a significant increase in neurodegeneration (>50%) when MnCl_2_ exposure was given after DA pretreatment. However, both adult and larval exposure to MnCl_2_ demonstrated significant toxicity by reducing the survival rate. In adult worms, 100 mM MnCl_2_ treatment after DA pretreatment further elevated the percentage of neurodegeneration. The Mn or DA alone exposed adult worms showed recovery of neuronal dopamine function within 24 hours, although exogenous DA and Mn treated worms showed prolonged behavioural defects. *Cat-2* mutants, without DA, were resistant to Mn mediated neurodegeneration. In contrast, *Cat-2* overexpressing strain displayed severe neurodegeneration at lower concentrations of MnCl_2_ (50 mM). Our results on biochemical, behavioural and genetic assays proved endogenous/exogenous DA level controls the sensitivity to Mn induced dopaminergic neurotoxicity.

## Introduction

In the central nervous system, functions are regulated through a series of neurotransmitters. Among them, dopamine plays a critical role in brain function, especially in behavioural modulations and movement [1–3]. Manganese (Mn) is an essential metal necessary for the efficient functioning of manganese-dependent enzymes and amino acid and lipid metabolism [4,5]. Chronic exposure to Mn may lead to manganism, which has symptoms that closely resemble idiopathic Parkinson’s Disease (iPD), including masked facial expression, gait impairment, and rigidity [6]. Mn exposed individuals have shown neuronal degeneration mainly in the dorsal striatum, internal globus pallidus, and substantial nigra pars reticulata (SNpr) – similar to Parkinson’s disease [7,8].

Elevated levels of iron (Fe) in SNpc regions of the brain and alterations in serum levels of Fe, manganese (Mn), copper (Cu), and zinc (Zn) in PD patients have been reported [9,10]. Both Fe and Mn share the same transporters. Due to nutritional deficiency, low levels of Fe can lead to the accumulation of Mn in brain regions such as the striatum, which might lead to cognitive deficits [11,12]. Excessive Mn is known to cause cell death by inducing oxidative stress and mitochondrial dysfunction. Recent studies also provide credence to the theory that environmental exposure to toxicants could initiate or propagate neurodegeneration by interfering with disease-associated proteins such as alpha-synuclein and amyloid proteins [13,14]. Hence, aberrancy in metal homeostasis could contribute to the pathophysiology of diseases like PD. Exposure to Mn is not limited to occupational exposure, but dietary exposure to Mn is also rising now. Mn is added to gasoline to increase its octane rating, as a disinfectant in water, and as an additive for parental nutrition and infant supplies [15,16].

Dopaminergic neurodegeneration is a hallmark of Parkinson’s disease (PD), with its onset associated with genetic susceptibility and ageing [11,17]. It has been postulated that “dysfunction” of the dopamine circuit results in movement disorders in Mn exposure, whereas degeneration of dopaminergic neurons occurs in PD [6]. Though it is generally accepted that individuals exposed to Mn have a higher risk of developing PD [18], the link between metal toxicity and the development of the disease is still debated. The studies on the death of dopamine neurons have suggested the involvement of multiple factors, including mitochondrial dysfunction, inflammatory response to the neurons, oxidative stress, attenuation in gene transcription levels, and translational depression of tyrosine hydroxylase gene, an essential enzyme involved in dopamine synthesis from tyrosine [19,20]. A recent study has shown that Mn exposure to *C. elegans* larvae affects the adaptive learning ability at later stages of its development [21,22].

The involvement of dopamine in Mn mediated neurotoxicity is not well understood. The susceptibility of dopamine neurons to Mn exposure during early development and its influence in altering neuronal function and behaviour has not been studied. In this study, we have used *C. elegans* as a model system to evaluate the effect of DA in augmenting dopaminergic neuronal toxicity.

## Materials and Methods

### Caenorhabditis elegans strains

All strains were maintained as previously mentioned in Nematode Growth Medium (NGM) plates with OP50 *E. coli* bacteria as a food source [23]. The strains were maintained at 20°C. The following strains were used in this study: N2 Bristol wild type; BZ555 [[egIs1 dat-1::pGFP]; UA44 [bal11;Pdat-1::α-syn+Pdat-1::gfp] gifted by Dr. Randy Blakely, BY200 [dat-1GFP(vtIs1)V]; MAB300 [dat-1::GFP(vtIs1) V, smf-2(gk133)] X -gift from Michael Aschner; CB1112 [*Cat-2*(e112)II] Gift by Dr. Gert Jansen; UA57 [baIs4 [dat-1p::GFP + dat-1p::*CAT-2*] gifted by Dr. Kim A Cladwell. All strains, unless otherwise mentioned, were provided by the *Caenorhabditis elegans* Centre (CGC, Minnesota).

### Manganese chloride preparation

For studying the effect of Manganese on *C. elegans*, different concentrations of MnCl_2_ (Merck-CAS #: 13446-34-9) solutions were used. A 1 M stock of MnCl_2_ was prepared in H_2_O from which the different concentrations of working solutions were made.

### Effect of exposure to manganese at the adult stage

All strains studied were exposed to 50 mM and 100 mM MnCl_2_ for 1 hour in a 20 µl final volume. The MnCl_2_ solution was removed by transferring the animals to a tube containing 1 ml distilled water, and the worms were then transferred to a glass slide and covered with a coverslip. Worms were anaesthetised using sodium azide (25 mM) before taking images. Microscopic observation was done for observing neurodegeneration markers as described before.

### Survival Test

Survival after acute MnCl_2_ treatment was tested in both adults and larval exposure. Larvae exposed to Mn (0 to 100 mM) were transferred to an OP-50 plate, and approximate initial numbers were recorded. 72-hour post-exposure, the number of live worms were counted by gently touching the worms by the tip of a worm picker. For adult worms, 20-25 worms were exposed to 50 mM and 100 mM MnCl_2_ concentrations at day 1 adult stage. After 1-hr MnCl_2_ exposure at 20°C, the worms were washed thrice in distilled water and transferred to OP-50 plate, and the number of live worms was recorded after 24 hours.

### Dopamine pre-treatment and MnCl_2_ exposure

Worms were treated with a concentration of either 5mM or10 mM DA (prepared from 1 M stock-Sigma-Aldrich-CAS # 62-31-7) for 10 min and then washed three times with M9 before exposing them to different concentrations of MnCl_2_ (0 -100 mM). Both locomotory and fluorescence intensity analyses were carried out following the treatment.

### Effect of Manganese on the worm movement: Locomotory assay

Day 1 adult worms were taken from the synchronised population of each strain and then exposed to Mn for 30 min at 20°C. The MnCl_2_ solution was removed by washing the worms thrice with distilled water before transferring them onto NGM plates with or without OP50. The movement of worms on both plates was then recorded using a Dino-Lite digital microscope. The worm locomotion was analysed by measuring body bends made by each worm in 20 seconds, manually counted, and recorded [24].

### Fluorescence intensity analysis

The images were taken using an inverted microscope Olympus IX51 equipped with Rolera-XR CCD camera (Q Imaging system) with image acquisition software NIS Elements-Advanced Research (NIKON). *C. elegans* worms of the treated and untreated group were placed on a glass slide containing 25 mM sodium azide in M9 buffer; once the worms were showing no movement, placed a cover glass over the worms. Worm images were taken at λ_ex_/λ_em_ 460-490/520. Fiji open software was used for image analysis.

### Microscopy and Camera

A stereo microscope (Magnus Analytics, India) with 10X zooming and Olympus IX51 inverted microscope (Olympus Imaging, Center Valley, PA, USA) that works with image acquisition software NIS Elements-Advanced Research (NIKON) and Rolera XR monochrome camera (QImaging, Canada) was used. A Dino-Lite EDGE camera (model# AM4115T-GFBW) was used to record the worms’ locomotory behaviour.

### Statistical analysis

GraphPad Prism was used to perform statistical analysis (GraphPad Software Inc.). To understand the effect of MnCl_2_ on 4 different strains we used two-way ANOVA with Tukey’s multiple comparison test to compare the percentage of neurodegeneration and survival. We conducted one-way ANOVA with Dunnett’s multiple comparison test to understand the effect of MnCl_2_ on single strain by comparing the percentage of neurodegeneration and survival. To estimate the percentage of neurodegeneration in UA57 strain we performed one-way ANOVA with Tukey’s multiple comparison test. In the results, data are represented as mean + SEM as indicated. Significance was represented as follows *p<0.05, **p<0.01, ***P<0.001,****p<0.0001.

## Results

### Dopamine is essential for manganese induced neurodegeneration

Our previous work has shown that Mn exposure to larval worms resist DAergic neurodegeneration [21] since a nominal DA expression in early larval stages [25]. To confirm whether the presence of DA is critical for MnCl_2_ to induce neurodegeneration in CEP neurons, we treated the adult worms with MnCl_2_ in four different strains of worms expressing GFP under dat-1 promoter: BZ555, UA44, MAB300, and BY200 (Fig. 1A). All the strains showed significant changes (p<0.0001; n≥3 trials) in CEP neurons after 100 mM MnCl_2_ treatment (Fig. 1A). Moreover, 50 mM MnCl_2_ treatment showed that MAB300 and BY200 are more sensitive (>20% showing neuronal changes; p<0.001 and p<0.05) compared to the other strains used (Fig. 1A). This results strongly suggest adult worms are more vulnerable to neurodegeneration with MnCl_2_ due to higher endogenous DA levels in comparison with the larval worms. However, no. of survivors was significantly low in all the strains tested. 100 mM MnCl_2_ treatment had a mortality rate of 50-80% (p<0.01, p<0.05, and p<0.01, for BZ555, UA44 and BY200 respectively) compared to the 20-40% in 50 mM (p<0.001, p<0.001, and p<0.01 for BZ555, UA44 and MAB300 respectively) MnCl_2_ treatment (Fig. 1B), suggesting a dose-dependent effect of MnCl_2_ in dopaminergic neuronal damage and survival.

**Figure 1:**
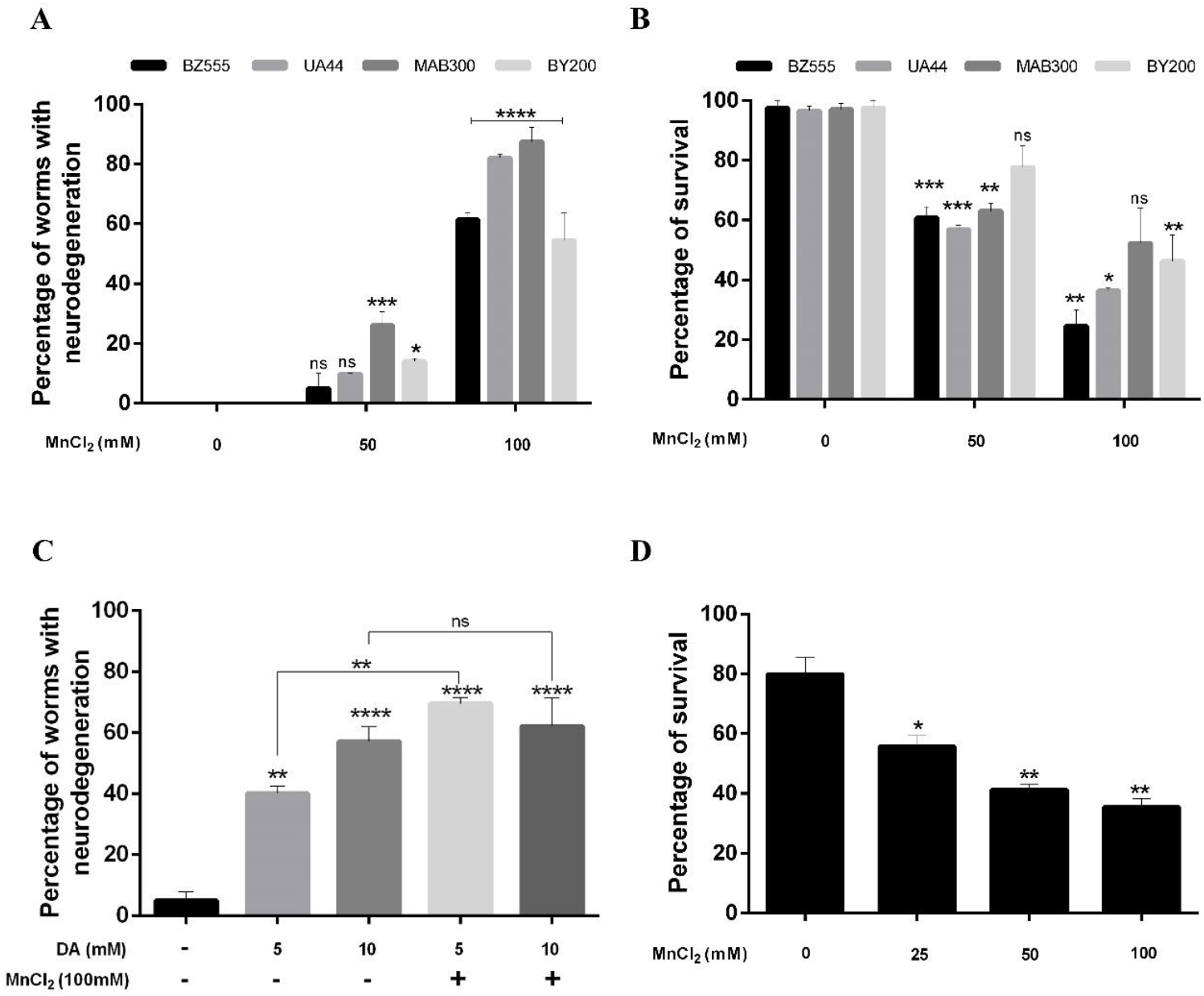
Larval exposure to MnCl_2_ shows resistance to neurodegeneration. **A**. Adult worms of BZ555, UA44, MAB300 and BY200 exposed to 50 mM and 100 mM concentrations of MnCl_2_ for 1-hour show percentage of neurodegeneration. n≥3 trials; each trial contains more than 15 worms; statistical analysis was carried out using two-way ANOVA with Tukey’s multiple comparisons test. **B**. Adult worms exposed to 50 mM and 100 mM MnCl_2_ for 1 hour showed a concentration-dependent reduction in survival. n≥3 trials; each trial contains more than 30 worms; statistical analysis was carried out using two-way ANOVA with Tukey’s multiple comparisons test. **C**. BZ555 larval worms pre-exposed to DA (both 5 mM and 10 mM) followed by 100 mM MnCl_2_ show neurodegeneration. n≥3 trials, each trial contains more than 15 worms; statistical analysis was carried out using one-way ANOVA with Dunnett’s multiple comparisons test. **D**. BZ555 larval worms exposed to increasing concentration MnCl_2_ show a significant concentration-dependent reduction in survival. n≥2 trials; each trial contains more than 100 worms; statistical analysis was carried out using one-way ANOVA with Dunnett’s multiple comparisons test. Data are represented as the mean +S.E.M. Significance indicated: ns-non-significant, p<*0.05, p< **0.01, p<***0.001 and p<***0.001.

If a higher level of DA expression in adults aids the Mn mediated neurodegeneration, we checked the effect of exogenous DA in larvae. Surprisingly, when the larval worms were exposed to MnCl_2_ after 5 mM DA pretreatment has shown an increase neurodegeneration; >60% neurodegeneration compared to 40% (p<0.01) in 5 mM DA alone (Fig. 1C). 10 mM DA treatment alone resulted in more than 50% (p<0.05; n≥3 trials) of worms showing neurodegeneration. Although, the addition of Mn did not show a significant (p<0.01) enhancing effect after 10 mM DA pretreatment compared to 10 mM DA alone (Fig.1C), suggesting that the treatment concentration might have reached the saturating point [26].

However, there was a significant reduction in the survival rate of larval worms exposed to MnCl_2_. Treatment at 25 mM, 50 mM and 100 mM concentrations of MnCl_2_ significantly (p<0.05, p<0.01 and p<0.01 respectively; n≥2 trials) reduced the survival rate from 60% (25 mM) to less than 40% (100 mM) (Fig. 1D). Treatment at higher concentrations (150 mM and 250 mM) was toxic and resulted in only a few survivors. This result show that excess level of endogenous DA itself is making the dopaminergic neuron vulnerable to neurodegeneration, and it exacerbate this effect in the presence of MnCl_2._

### Dopamine neurotoxicity in adults enhanced in the presence of exogenous dopamine

Next, we wanted to test the effect of exogenous DA on adult worms in the presence and absence of MnCl_2_. DA pretreatment in BZ555 adult worms showed dopaminergic neuronal damage on CEP and ADE neurons when presented alone (Fig 2B) in comparison with the control (Fig 2A). Additionally, MnCl_2_ exposed after 10 mM DA pretreatment showed severe neurodegeneration markers such as puncta, blebbing’s and breaks at 50 mM (Fig 2C) and in 100 mM MnCl_2_ (Fig 2D) along with CEP loss. Our result strongly suggests exogenous DA is crucial for potentiating severe neurodegeneration in response to MnCl_2_ on dopaminergic neurons. Adult worms exposed to MnCl_2_ alone did not show such type of neurodegeneration pattern [21].

**Figure 2.**
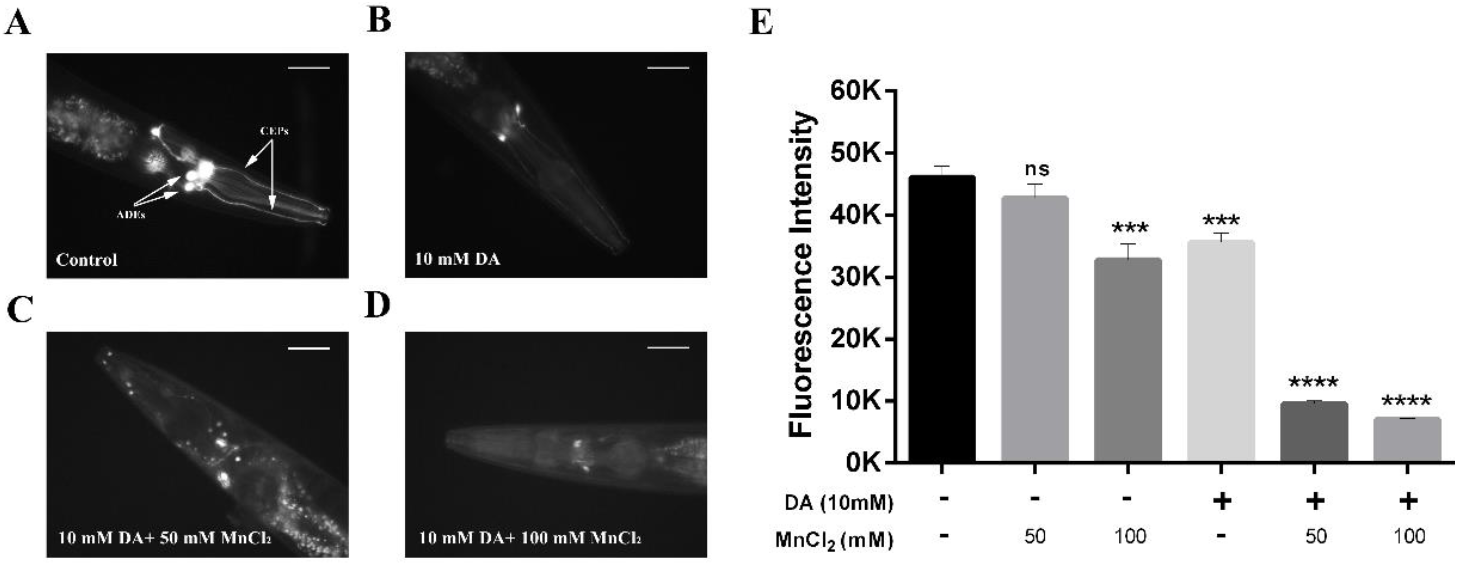
Adult worms were showing DAergic neurodegeneration in response to MnCl_2_ and DA exposure. **A-D**. Representative images of BZ555 CEP and ADE dopaminergic neurons pre-exposed to DA for 10 min, followed by 1-hour MnCl_2_ exposure. **A**. Control worms show continuous expression of GFP on CEP and ADE. **B**. Worms pre-exposed to 10 mM DA show reduction in GFP and size of ADE neurons as a marker of neurodegeneration. **C**. 50 mM, MnCl_2_ exposure to worms pre-exposed with DA show neurodegeneration markers such as puncta, breaks on CEP neurons in addition to the loss of soma in ADE neurons. **D**. 100 mM MnCl_2_ exposure to worms pre-exposed with DA show severe neurodegeneration such as loss of CEP neurons and reduced GFP expression on ADE neurons along with shrinkage. n≥3 trials; each trial contains more than 15 worms; scale bar represents 50 µm, exposure 500 ms. **E**. Quantification of the dat-1::gfp level of day-1 BZ555 strain on exposure to MnCl_2_ and DA. n≥ 15 worms; statistical analysis was carried out using one-way ANOVA with Dunnett’s multiple comparisons test. Data are represented as the mean + S.E.M. Significance indicated: ns-non-significant, p<***0.001 and p<****0.0001).

Additionally, to understand the magnitude and cellular degeneration, we measured dat-1 protein expression levels of CEP and ADE neurons exposed to MnCl_2_ with or without DA pretreatment. The data showed a significant reduction in dat-1 expression levels in worms treated with 100 mM MnCl_2_ compared to the control (p<0.001), whereas 50 mM did not show a significant reduction in dat-1 expression (Fig 2E). Pre-treatment with 10 mM DA alone shows a significant reduction in dat-1 expression, is giving additional evidence for DA alone can cause severe DA neurodegeneration (Fig 2E). A similar result was found when DA was given to larval worms, supporting strong evidence for DA mediated DAergic neurodegeneration (Fig 1C). DA pretreatment and subsequent MnCl_2_ exposure (both 50 and 100 mM) resulted in significant reduction (p<0.0001; n≥15 worms) in dat-1 expression level (Fig. 2E). Altogether the result suggests that exogenous DA constitutes significant pathological changes in DAergic neurons in the presence and absence of MnCl_2_.

### Partial recovery from Mn toxicity over time with altered dopamine sensitivity

To know whether the worms can recover from Mn-induced toxicity, we tested on-food and off-food behaviour of wild-type, a characteristic test for the DA function in the worm post-exposure to Mn (Fig. 3A) [24]. The behaviour of no exposure control worms was recorded at different time points from 0-24 hours period (Fig. 3B). A significant (p<0.0001;n≥15 worms) slowing down behaviour of worms after 30 minutes of starvation was observed on the on-food plate (Fig. 3B). Worms from the same batch were tested after 30 min, 3 hours, and 24 hours. The on-food behaviour recovered from the initial extreme slow-down was maintained at a lower level than in the off-food; this is mainly due to the surge of DA in the presence of food, which results in significant slowing down behaviour of the worm. Under similar conditions in the control worms pre-treated with 10 mM DA, its off-food behaviour also showed a significant reduction (p<0.0001) in body bends at 0^th^ hour compared to no exposure control (Fig. 3C). After DA was washed off, the recovery from the DA plus starvation took around 24 hours to reach that of the control animal behaviour in on-food (Fig.3 C). Off-food behaviour recovered to that of control within 30 minutes after the removal of DA (Fig. 3C). This result suggests DA and Mn alone exposure affects the behaviour of the worms, plausibly resulting from the neurodegeneration as observed earlier (Fig. 1A and 1C).

**Figure 3.**
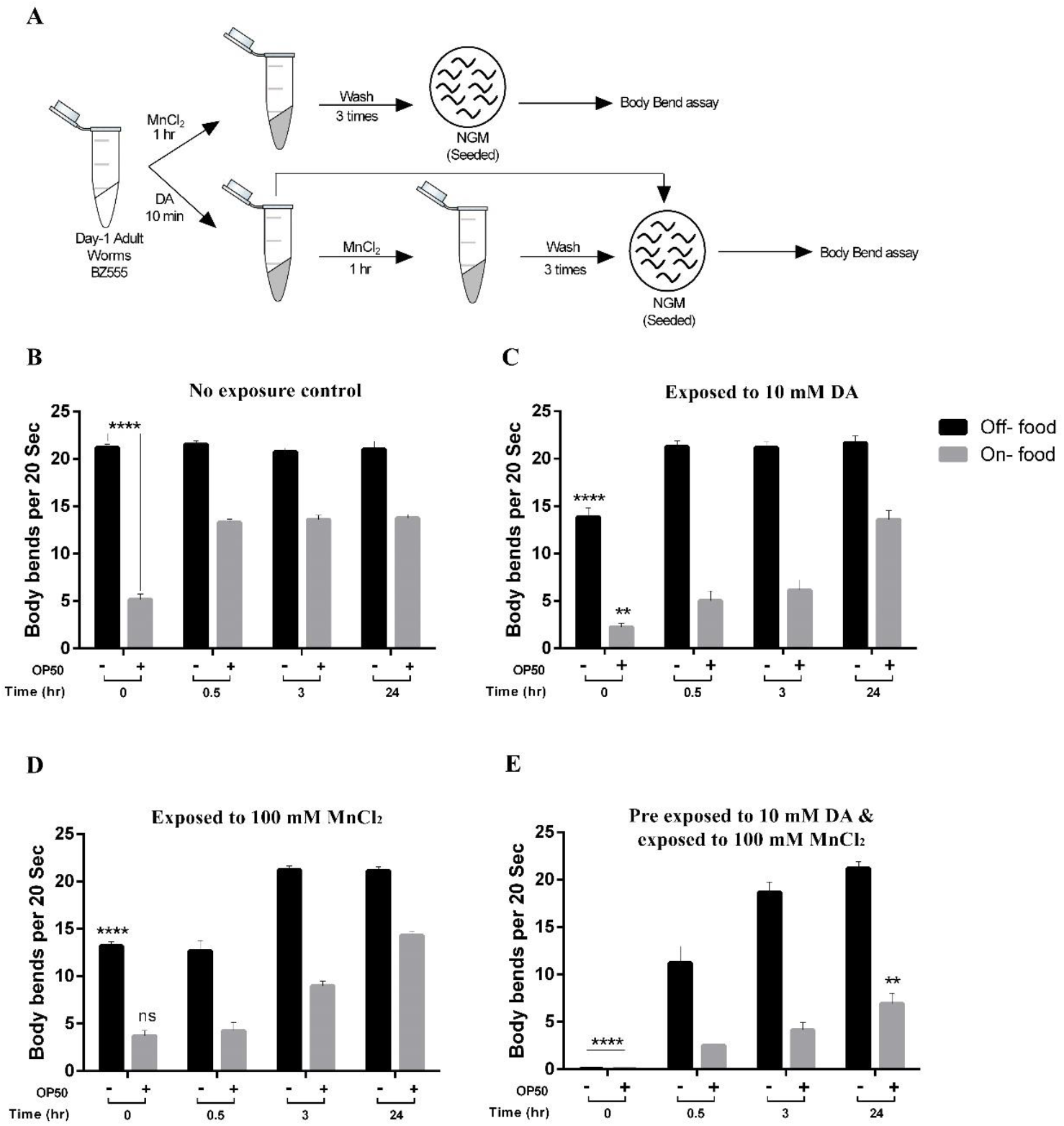
Recovery time kinetics of body bends in off-food and on-food. **A**. The protocol for exposure of BZ555 adult worms to DA and MnCl_2_. **B**. Off-food and on-food body bend behaviour of no exposure control. **C**. Off-food and on-food body bend behaviour of DA pre-exposed adult worms. **D**. Off-food and on-food body bend behaviour of MnCl_2_ exposed adult worms. **E**. Off-food and on-food body bend behaviour of worms pre-exposed to DA followed by MnCl_2_. n≥15 for all four sets of experiments. All statistical analysis was carried out using two-way ANOVA with Tukey’s multiple comparisons test. Data are represented as the mean + S.E.M. Significance indicated: ns-non-significant, p< *0.05, p< **0.01, p<***0.001 and p<****0.0001).

However, 100 mM MnCl_2_ exposure showed significant reduction (p<0.0001 and p<0.01 respectively) in body bends at 0^th^ hour compared to no exposure control off-food and on-food body bends, the off-food plate’s recovery took less than 3 hours, and the on-food behaviour took around 24 hours (Fig. 3D). Pretreatment of DA and subsequent MnCl_2_ exposure completely immobilized the worms at 0^th^ hour (p<0.0001) in both on-food and off-food plates (Fig. 3E). These worms showed a slower recovery in off food plate (in less than 3 hours) and completely recovered before 24 hours. However, the on-food behaviour showed significantly lower (p<0.01) recovery than the no exposure control even after 24 hours (Fig. 3E), further suggesting that the sensitivity to MnCl_2_ mediated toxicity in the presence of DA has increased in these worms.

### The absence of dopamine leads to resistance against MnCl_2_ mediated neurodegeneration

We wanted to test the role of DA in MnCl_2_ induced toxicity in further detail. Instead of exogenous addition of DA, we used two strains with altered endogenous DA levels: CB1112 strain, deficient in tyrosine hydroxylase gene (*cat-2*) with low DA synthesis, and UA57 having an extra copy of *cat-2* gene, with an increased level of DA synthesis. CB1112 showed slightly higher body bends (>12%; p>0.05;n≥15 worms) compared with wild type without any treatment (Fig. 4A). On the other hand, UA57 showed a moderate reduction in the number of body bends (<22%, p<0.05) compared to the wild-type worms (WT) (Fig. 4A). When CB1112 worms were exposed to DA, the body bend behaviour did not alter significantly but maintained a substantially higher percentage of body bends (>57%; p<0.01) compared to that of WT (Fig. 4A). This behaviour of CB1112 could be possibly due to the compensatory effect of exogenous DA in the absence of the functional *cat-2* gene in this worm. When UA57 worms were treated with MnCl_2_, it further lowered the body bend behaviour to >82% (p<0.0001) compared to the WT treated with MnCl_2_ (Fig. 4A). CB1112 strain, on the other hand, did not show any reduction in body bends in the presence of MnCl_2_ compared to the WT (Fig. 4A). This result further confirms the role of intracellular DA content in MnCl_2_ mediated dopaminergic neuronal degeneration. DA and MnCl_2_ alone treatment did not affect the body bend behaviour of CB1112 but rather showed increased body bends (57 and 63% respectively; p<0.01) compared to that of their respective WT control. However, when pre-treated with DA and subsequently exposed to MnCl_2_, CB1112 worms showed a significant reduction in body bends (>29%; p>0.05) compared to the WT (Fig. 4A). Altogether, the results revealed that CB1112 worms with a deficient level of DA show resistance to Mn treatment with normal locomotory behaviour, while UA57 with higher internal DA level show higher sensitivity to MnCl_2_ mediated neurotoxicity by completely reduced locomotory behaviour.

**Figure 4.**
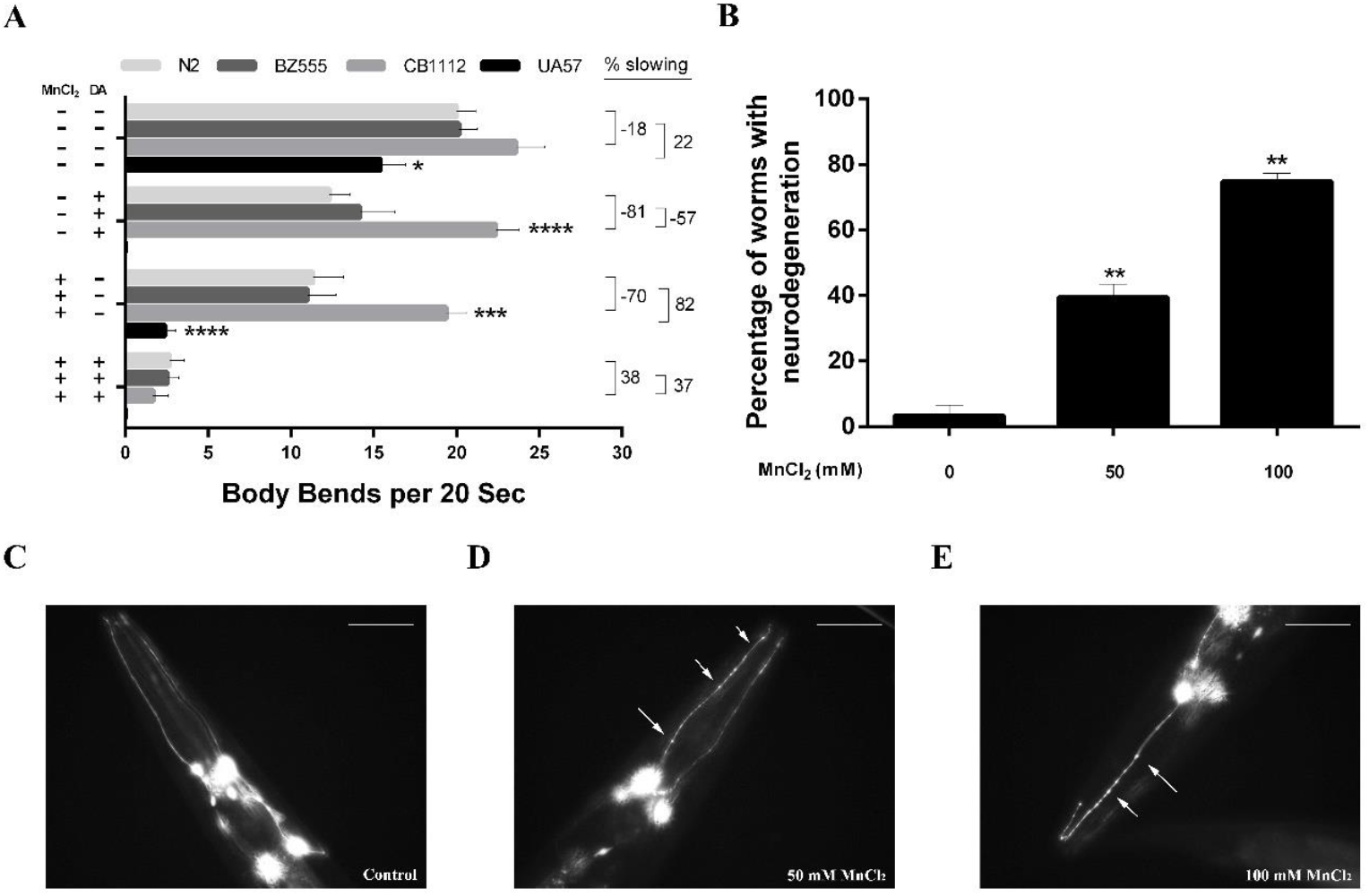
Dopamine level determines neurodegeneration in UA57 and CB1112 strain. **A**. Off-food behaviour of the DA synthesis enzyme, tyrosine hydroxylase gene (*cat-2*) mutants. No exposure behaviour of both strains was assessed along with N2 (WT) and BZ555 as a control for genetically modified strains. n≥15 worms; statistical analysis was carried out using two-way ANOVA with Tukey’s multiple comparisons test. **B**. *cat-2* overexpressing UA57 adult worms exposed to MnCl_2_ alone show a significant increase in the percentage of neurodegeneration at both concentrations of 50 mM and 100 mM MnCl_2_. n≥2 trials; each trial contains more than 20 worms; statistical analysis was carried out using one-way ANOVA with Tukey’s multiple comparisons test. **C-E**. Representative images of *cat-2* overexpressing UA57 worms show neurodegeneration markers on exposure to MnCl_2_, such as puncta and blebbing (50 mM MnCl_2_) and breaks and loss of CEP neurons (100 mM MnCl_2_) along with other neurodegeneration signs (white arrows). n≥3 trials; each trial contains more than 15 worms; scale bar represents 50 µm, exposure 500ms. Data are represented as the mean + S.E.M. Significance indicated: ns-non-significant, p< *0.05, p< **0.01 and p<****0.0001).

### Overexpression of cat-2 gene enhances Mn dependent DAergic neurodegeneration

In addition to the reduction of locomotory behaviour, we wanted to observe how DA neurons in *cat-2* overexpressing UA57 strain responds to MnCl_2_. UA57 adult worms showed a significant level (>40%; p<0.01) of neurodegeneration at 50 mM MnCl_2_ exposed (Fig 4B). In contrast, at 100 mM MnCl_2_ exposure, the UA57 strain showed a significantly greater percentage (>75%, p<0.01) of neurodegeneration compared to the control (Fig. 4B). Neurodegeneration observed in UA57 at 50 mM MnCl_2_ was significantly higher than the other strains BZ555, UA44 and BY200 at 50 mM MnCl_2_ exposed (p<0.0001, p<0.001 and p<0.01 respectively) (see Fig. 1A and 4B). Moreover, BZ555 and BY200 (p<0.01) strains were showed a lesser degree of degeneration compared to UA57 at 100 mM (see Fig. 1A and 4B). These results suggest that a higher DA level inside the worms is crucial to determining Mn mediated DA neurodegeneration.

## Discussion

Animal behaviour is a combinatorial effect of the information processed by the nervous system based on the external stimuli. DA is a critical neurotransmitter that drives the fundamental behaviours associated with movement, reward, attention, and cognition [27,28]. Perturbations in DA levels results in behavioural disorders such as addiction, ADHD, Schizophrenia and Parkinson’s disease [29,30].

Our previous study has shown that larval worms are resistant to Mn mediated DA neurodegeneration [21]. In this study, adult worms exposed to Mn has shown significant neurodegeneration characterised by markers such as puncta, blebbing’s, breaks and CEP loss. This result indicates that the stage-specific expression of tyrosine hydroxylase (TH), encoded by *cat-2* gene, an enzyme required to synthesise DA, promoted Mn-mediated neurodegeneration in adult worms [31,32,25]. In contrast, the resistance to Mn mediated neurotoxicity at the larval stage was diminished in the presence of exogenous DA, which implies that the DA synthesis was poor at early stages. Mn exposure without exogenous DA at the L1 stage showed no neurodegeneration further proving the stage specific expression of DA has vital role in Mn mediated neurodegeneration. It has been noted that endogenous DA enhances mitochondrial respiratory inhibition and free radical production in the striatum [33].

Altogether, in the present study, the presence of DA, endogenous in adults and exogenous in the larval worm, is critical in determining the potential of Mn directed neurodegeneration.

L-DOPA (3,4-dihydroxyphenylalanine) is the most effective drug prescribed for patients with Parkinson’s disease for its symptomatic treatment clinically [34]. However, in-vitro and pre-clinical studies demonstrate L-DOPA toxicity on dopaminergic neurons are accelerated by its derivatives through oxidative damage [34,35]. In rat models, combined infusion of MPTP and L-DOPA generated a significant reduction of nigral neurons reasonable to suggest that L-DOPA could potentially produce dopaminergic neurotoxicity [36]. Neuromelanin formed from DA oxidation leads to the generation of many o-quinones, which guides mitochondrial dysfunction, α-synuclein aggregation into neurotoxic oligomers, and impaired protein degradation [37]. In DJ-1 knockout and A53T α-synuclein transgenic mice, DA content is needed to show PD’s pathological features [38].

Correspondingly, our results on adult worms showing exogenous DA content advances DAergic neurodegeneration in the presence of Mn provide evidence for DA action in PD pathogenesis in vivo.

DA is one of the major neurotransmitters modulating the locomotory behaviour in the presence and absence of food [24]. Additionally, during the L1 stage, exogenous DA altered its on-food/off-food behaviour, almost mimicking the effects of Mn exposure. Both Mn and DA exposure at the L1 stage has a long-term impact on the on-food/off-food animal behaviour [21]. These worms developed to the adult stage with characteristic signs of neuronal damage in dopaminergic neurons. In mice, it has been shown that Mn exposure during embryonic stages affects neurogenesis and shows a higher number of immature GABAnergic neurons in the dentate gyrus [39]. An extensive set of genes was found to downregulate the transcripts because Mn induced hypermethylation at the promoter region [40]. Proper DNA methylation, histone modifications, and non-coding RNA (mainly miRNA and NRSE dsRNA) expressions are essential regulators of neurogenesis and connectome formations [41].

DA release at the synapse is essential to induce Mn associated neurodegeneration. Dopaminergic cells are prone to direct toxicity of Mn^2+^ or Mn^3+^ through induction of reactive oxygen species (ROS) [42]. Mn^2+^ induces catabolism of dopamine to 6-hydroxydopamine or other toxic catecholamines and decreases the antioxidant like GSH, thiols etc. [43]. Mn^3+^ ions and their pyrophosphates can rapidly oxidise dopamine and generate semiquinones and orthoquinone by sequential oxidation [44]. A continuous redox cycling between Mn^2+^ and Mn^3+^ is also believed to be occurring, resulting in DA oxidation and generating aminochrome [45], which induces acute cell death [46]. Our results on larval and adult worms showed that exogenous/endogenous DA is essential to cause severe neurodegeneration results in loss of developmental, physiological and behavioural functions. *Cat-2* mutants showed resistance to Mn induced neurotoxicity, while the *Cat-2* overexpressing UA54 strain showed significant neurotoxicity to dopaminergic neurons on Mn exposure. These results confirm the earlier observation that both endogenous and exogenous DA is directly involved in the Mn induced neurotoxicity [47].

In conclusion, this study provides new insights about the functional role of DA dependant enhancement of Mn induced DAergic neurodegeneration. Both exogenous and endogenous DA studies emphasise the vulnerability of DA neurons to Mn exposure and show possible modes of the pathogenesis of PD. Further understanding of these changes would help decipher the disease onset both in Manganism and PD, with a view of a better treatment strategy.

## Acknowledgement

We thank Dr Micahel Aschner-Albert Einstein College of Medicine, Dr Kim. A. Cladwell-University of Alabama, Dr Gert Jansen-University of Nijmegen and Dr Randy Blakely-Florida Atlantic University and CGC, Minnesota for providing us with some of the strains used in the study. We thank Ms Rasitha Santhosh Kanakalatha, Ms Agrima Nair, Ms Aswathy AR, Mr Amal Wilson Varghese and Ms Swathy S Nair for their active participation in improving the work. A fellowship from SCTIMST supported VR. This work was partially supported by the TRC grant (AT).

## Conflict of Interest

The authors declare that they have no conflicts of interest.

## Notes

### Competing Interest Statement

The authors have declared no competing interest.

